# Synthetic dual-function RNA reveals features necessary for target regulation

**DOI:** 10.1101/2021.06.30.450655

**Authors:** Jordan J. Aoyama, Medha Raina, Gisela Storz

**Affiliations:** Division of Molecular and Cellular Biology, Eunice Kennedy Shriver National Institute of Child Health and Human Development, Bethesda, MD 20892-5430, USA

**Keywords:** MgrR, MgtS, Hfq, sRNA, small protein, synthetic biology

## Abstract

Small base pairing RNAs (sRNAs) and small proteins comprise two classes of regulators that allow bacterial cells to adapt to a wide variety of growth conditions. A limited number of transcripts encoding both of these activities, regulation of mRNA expression by base pairing and a small regulatory protein, have been identified. Given that few have been characterized, little is known about the interplay between the two regulatory functions. To investigate the competition between the two activities, we constructed synthetic dual-function RNAs, hereafter referred to as MgtSR or MgtRS, comprised of the *Escherichia coli* sRNA MgrR and the open reading frame encoding the small protein MgtS. MgrR is a 98 nt base pairing sRNA that negatively regulates *eptB* encoding phosphoethanolamine transferase. MgtS is a 31 aa small inner membrane protein that is required for the accumulation of MgtA, a magnesium (Mg^2+^) importer. Expression of the separate genes encoding MgrR and MgtS is normally induced in response to low Mg^2+^ by the PhoQP two-component system. By generating various versions of this synthetic dual-function RNA, we probed how the organization of components and the distance between the coding and base pairing sequences contribute to the proper function of both activities of a dual-function RNA. By understanding the features of natural and synthetic dual-function RNAs, future synthetic molecules can be constructed to maximize their regulatory impact.

**IMPORTANCE:** Dual-function RNAs encode a small protein and also base pair with mRNAs to act as small, regulatory RNAs. Given that only a limited number of dual-function RNAs have been characterized, further study of these regulators is needed to increase understanding of their features. This study demonstrates that a functional synthetic dual-regulator can be constructed from separate components and used to study the functional organization of dual-function RNAs, with the goal of exploiting these regulators.

## INTRODUCTION

Small RNAs (sRNAs) play an important role in shaping and regulating cellular responses to changing environmental conditions (reviewed in (1)). Hundreds of sRNAs have been identified through various deep sequencing metholodogies (reviewed in (2, 3)). Most of these 50-300 nucleotide (nt) molecules regulate target mRNAs through base pairing (reviewed in (4)). In many bacteria, sRNAs depend on the RNA binding protein Hfq (<u>h</u>ost <u>f</u>actor for bacteriophage <u>Q</u>β) for stability and efficient base pairing with their target mRNAs (reviewed in (5, 6)). The functional outcomes of sRNA base pairing with the target mRNA include the inhibition of translation, an increase in translation, a change in mRNA degradation, as well as post-initiation decreases in transcription (reviewed in (7, 8)).

While generally assumed to be noncoding, some sRNAs have been found to encode small ORFs (sORFs) and for a small subset of the RNAs, this ORF has been shown to be translated. An sRNA capable of base pairing regulation while also encoding a protein is denoted a dual-function RNA (reviewed in (9)). Until now, only a limited number of dual-function RNAs have been identified and characterized as they tend to be overlooked due to the difficulty in distinguishing true translated ORFs from sORFs that occur randomly. Furthermore, traditional biochemical methods for identifying and purifying proteins commonly miss proteins that are less than 50 amino acids (reviewed in (10)).

Despite the difficulties inherent in the study of dual-function RNAs, two *E. coli* dual-function RNAs, SgrST and AzuCR, have been examined in sufficient depth to describe both the base pairing and protein coding activities. Both of these RNAs are involved in regulating carbon metabolism. The 227 nt SgrS sRNA was originally identified in a computational screen for sRNAs in *E. coli* (11). Subsequently, SgrS was shown to play a role in the cellular response to glucose-phosphate stress and was found to base pair with the *ptsG* and *manXYZ* mRNAs, which encode important sugar transporters (12). In the course of these studies, it was discovered that SgrS is translated to produce the 43 amino acid small protein SgrT, which inhibits the transport activity of the glucose permease PtsG (13). Together, SgrS and SgrT counteract the accumulation of sugar phosphates in the cell by reducing the translation and activity of sugar transporters and increasing dephosphorylation of sugar phosphates. Another recently characterized dual-function RNA is AzuCR (14). This 164 nt RNA was initially identified in a bioinformatic search for novel sRNA genes but was reclassified as an mRNA when it was found to encode a 28 aa protein, AzuC (15, 16). AzuC interacts with aerobic glycerol-3-phosphate dehydrogenase (GlpD) to increase its activity (14). However, overexpression of AzuCR produces growth defects in galactose and glycerol medium that persist even when an AzuCR derivative carrying a stop codon mutation in the sORF is overexpressed, suggesting that AzuCR also functions as an sRNA. Consistent with this observation, AzuCR is capable of direct base pairing regulation of *cadA*, encoding lysine decarboxylase, and *galE*, encoding UDP-glucose 4-epimerase.

One interesting question regarding dual-function RNAs is whether the two functions can be carried out simultaneously by the same molecule or whether the activities interfere with each other. The base pairing region of SgrS is 15 nt downstream of the SgrT ORF. Mutations to inhibit *sgrT* translation do not impede SgrS regulation of mRNA targets, but mutations that restrict base-pairing increase SgrT translation indicating that the base-pairing function interferes with translation. Furthermore, translation of the small protein lags behind transcription of SgrS RNA by approximately 30 min (17). Based on these observations, it was suggested that the base pairing function predominates, and SgrST molecules that base pair with target mRNAs are unavailable to be translated as they are co-degraded. Once the pool of targeted mRNAs has been depleted, SgrST accumulates and is translated.

In contrast, there is direct overlap between the base pairing region and sORF of AzuCR such that one function necessarily interferes with the other (14). The introduction of a stop codon, which presumably reduces ribosomal occupancy of the transcript increases base pairing activity. For AzuCR, translation of the small protein is repressed by the anaerobic-growth induced sRNA FnrS suggesting sRNAs are one mechanism by which the competition between the base pairing and protein coding activites are controlled.

To further explore the features of dual-function RNAs that impact what activity predominates, we chose to construct synthetic dual-function RNAs. We decided to use parts of the individual transcripts corresponding to the sRNA MgrR and encoding the small protein MgtS. The sRNA component, MgrR, was identified through Hfq co-immunoprecipitation followed by genome-wide RNA detection on microarrays (18). This sRNA negatively regulates *eptB*, a lipopolysaccharide (LPS) modifiying enzyme, and *ygdQ*, a protein of unknown function (19). MgtS is a 31 amino acid small protein that interacts with MgtA, a P-type ATPase Mg^2+^ importer (20). This small protein was first predicted as a conserved sORF and further studies showed that it is translated in *E. coli* (11, 21). The interaction of MgtS with MgtA serves to increase levels of the importer and increase the intracellular concentration of Mg^2+^. Induction of both components, MgrR and MgtS, is dependent on the two-component system, PhoQ/PhoP. In low Mg^2+^ conditions, the sensor kinase PhoQ autophosphorylates and donates a phosphate group to PhoP. Phosphorylated PhoP in turn activates a number of different genes including *mgrR* and *mgtS*, which are needed for survival in low Mg^2+^ (22).

In this study, we demonstrate that it is possible to create a synthetic dual-function RNA from existing individual components, in this case from the transcripts encoding MgrR and MgtS. Additionally, we show that the composition of the sequence between the functional elements impact their activities.

## RESULTS

### Construction of a synthetic dual-function RNA

We set out to build a synthetic dual-function RNA from two components that could be easily tested for their regulatory activities. We chose MgrR as the sRNA component and MgtS as the small protein component. The corresponding two genes are transcribed convergently in a PhoPQ-dependent manner in response to low Mg^2+^ conditions (Fig. 1A). The design of the first synthetic RNA denoted MgtSR was modeled after the characterized dual-function RNA, SgrST, whose sORF precedes the base pairing region by 15 nt (23). Our design included the MgtS ORF, from the start codon to the stop codon (96 nt), immediately followed by 77 nt of native *mgrR* sequence (see Fig. S1 in supplemental material). The pBAD24 plasmid carrying the construct provided the ribosome binding site (RBS) for MgtS. The sequence from *mgrR* includes a 9 nt Hfq A-R-N (where A = adenine, R = adenine or guanine, N = any nucleotide) binding motif, the seed sequence for base pairing interactions with MgrR targets, and an intrinsic terminator with a polyU stretch also bound by Hfq (5, 24). The MgtS ORF and the seed sequence of MgrR are separated by 30 nt of native sequence from MgrR (denoted a spacer here) (Fig. 1B). The hybrid gene was cloned behind an arabinose-inducible promoter to control expression.

**FIG. 1.**
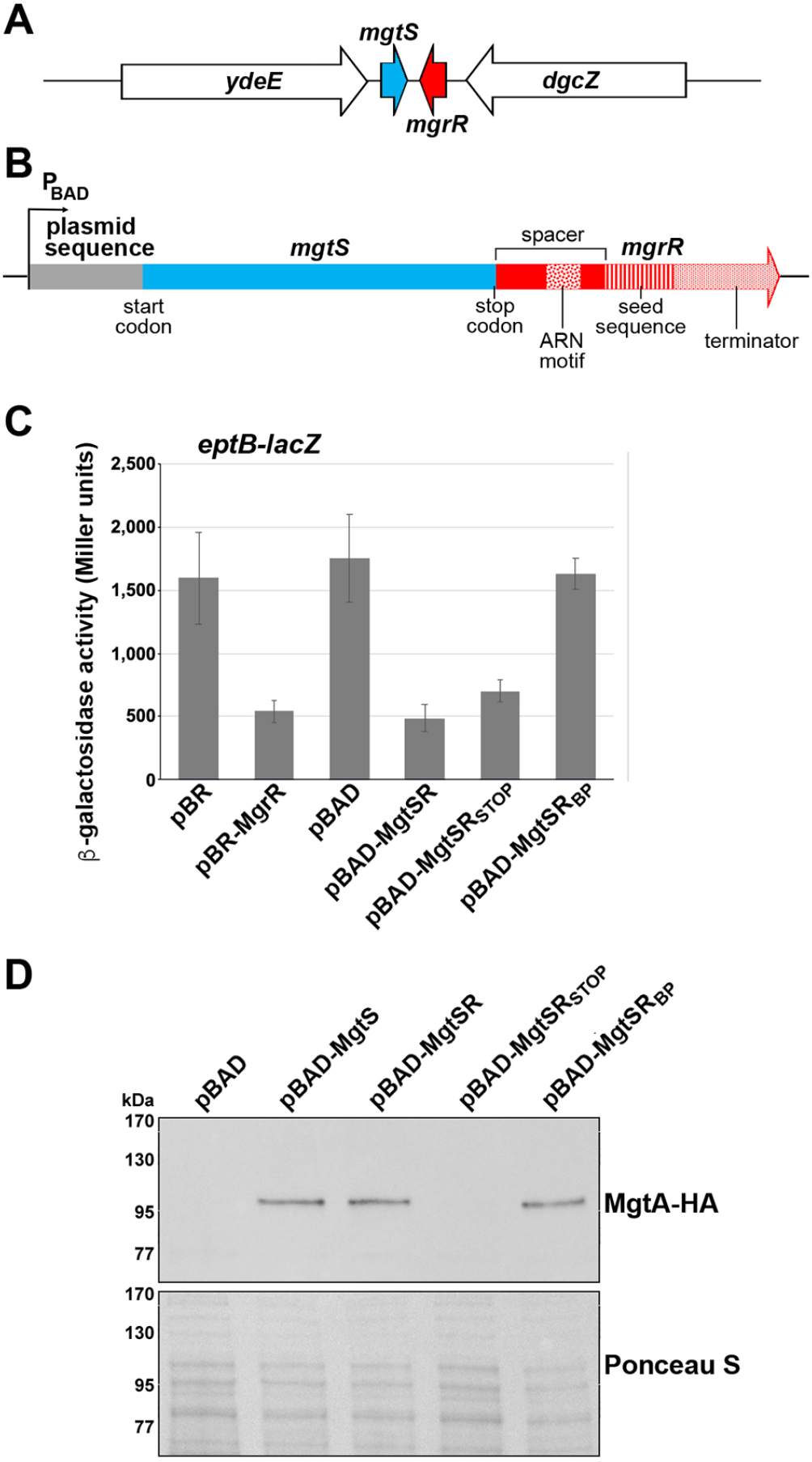
Regulatory activity of a synthetic dual-function RNA. (A) Diagram of the endogenous location of *mgtS* (blue) and *mgrR* (red). (B) Schematic of the hybrid dual-function RNA MgtSR. Plasmid sequences are in gray, *mgtS* sequences are in blue and *mgrR* sequences are in red. The ARN, seed, and terminator sequences are indicated by distinct patterns as labeled. (C) Effect of pBR, pBR-MgrR, pBAD, pBAD-MgtSR, pBAD-MgtSR_STOP_, or pBAD-MgtSR_BP_ overexpression on an *eptB-lacZ* fusion in Δ*mgrR* background. Graphs show average of three biological replicates with error bars representing standard deviation. (D) Immunoblot analysis of MgtA-HA cells in a Δ*mgtS* background transformed with pBAD, pBAD-MgtS, pBAD-MgtSR, pBAD-MgtSR_STOP_, or pBAD-MgtSR_BP_ grown to OD_600_ 0.4 in N-media supplemented with 500 μM Mg^2+^, washed, and then resuspended and grown in N-media supplemented with arabinose but without Mg^2+^. Samples were collected after 20 min, and MgtA-HA was visualized with α-HA antibodies to detect MgtA-HA. The Ponceau S provides the loading control.

We also designed controls to individually eliminate the activity of each component. To this end we constructed a stop codon control, pBAD-MgtSR_STOP_, to prevent translation of the small protein component while maintaining the integrity of the base pairing region, and pBAD-MgtSR_BP_, a construct with mutations in the base pairing region of MgrR to prevent sRNA activity while maintaining translation of the small protein.

### MgtSR is functional

To determine whether MgtS is functional, we first assayed the effect of MgtSR overexpression on the activity of an *eptB*-*lacZ* fusion, a reporter fusion to a known target of MgrR. β-galactosidase activity was measured in Δ*mgrR* cells carrying the *eptB*-*lacZ* translational fusion and transformed with pBR, pBR-MgrR, pBAD, pBAD-MgtSR, pBAD-MgtSR_STOP_, and pBAD-MgtSR_BP_. Repression of *eptB*-*lacZ* was observed upon overexpression of MgtSR to a similar extent as overexpression of MgrR (Fig. 1C). Furthermore, pBAD-MgtSR_STOP_ derivative was still able to repress the fusion, but pBAD-MgtSR_BP_ no longer affected *eptB-lacZ* expression. This result demonstrates that the MgtSR RNA has the ability to act as a base pairing regulatory RNA.

We similarly tested for functional expression of the MgtS component by carrying out immunoblot analysis to examine the levels of MgtA, a membrane protein that is stabilized by MgtS (20). Δ*mgtS* cells expressing chromosomally-encoded HA-tagged MgtA were transformed with pBAD, pBAD-MgtS, pBAD-MgtSR, pBAD-MgtSR_STOP_, and pBAD-MgtSR_BP_. Overexpression of MgtSR increases MgtA-HA levels under conditions of Mg^2+^ limitation similar to MgtS (Fig. 1D). As expected, pBAD-MgtSR_STOP_ did not affect MgtA-HA levels while pBAD-MgtSR_BP_ did, demonstrating that a functional small protein is translated from *mgtSR_BP_* despite the inability of the transcript to repress *eptB-lacZ* (Fig. 1D). The β-galactosidase data taken together with the immunoblot analysis indicate that we have successfully constructed a synthetic dual-function RNA that can act as a base pairing sRNA and also express a functional small protein.

### Translation of *mgtS* is dominant in the absence of a spacer

To examine how the *mgrR* sequence that provides a spacer between the *mgtS* coding sequence and the *mgrR* base pairing region impacts the activities of the two MgtSR components, we removed the 30 nt sequence to generate the pBAD-MgtSR-no spacer construct (Fig. 2A). We hypothesized that in the absence of the spacer, the two functions may interfere with each other resulting in the loss of one or both of the activities. To test this hypothesis, we asssayed the *eptB-lacZ* fusion and MgtA-HA levels for cells transformed with pBR, pBR-MgrR, pBAD, pBAD-MgtSR, and pBAD-MgtSR-no spacer. We discovered that removing the spacer sequence between the components of MgtSR eliminates the regulation of the *eptB*-*lacZ* fusion (Fig. 2B) but not stabilization of MgtA (Fig. 2C). These results suggest that translation of MgtS predominates and may sterically block MgrSR base pairing activity (25). To test this prediction, we introduced a stop codon into the pBAD-MgtSR-no spacer construct. This construct, pBAD-MgtSR-no spacer_STOP_, rescued *eptB*-*lacZ* regulation consistent with translation of *mgtS* disrupting base pairing regulation by MgtSR (Fig. 2B). It is interesting to note, that the MgtSR-no spacer_STOP_ RNA is still functional despite the upstream Hfq binding site being removed, though sequencing in the *mgtS* coding sequence could compensate. We also noted that the levels of this sRNA are significantly higher than MgtSR (see Fig. S2A in supplemental material). As expected, the MgtSR-no spacer_STOP_ RNA, which should not be translated, did not lead to increased levels of MgtA-HA. Thus, the presence of a 30 nt spacer sequence helps to maintain the independent function of each component.

**FIG. 2.**
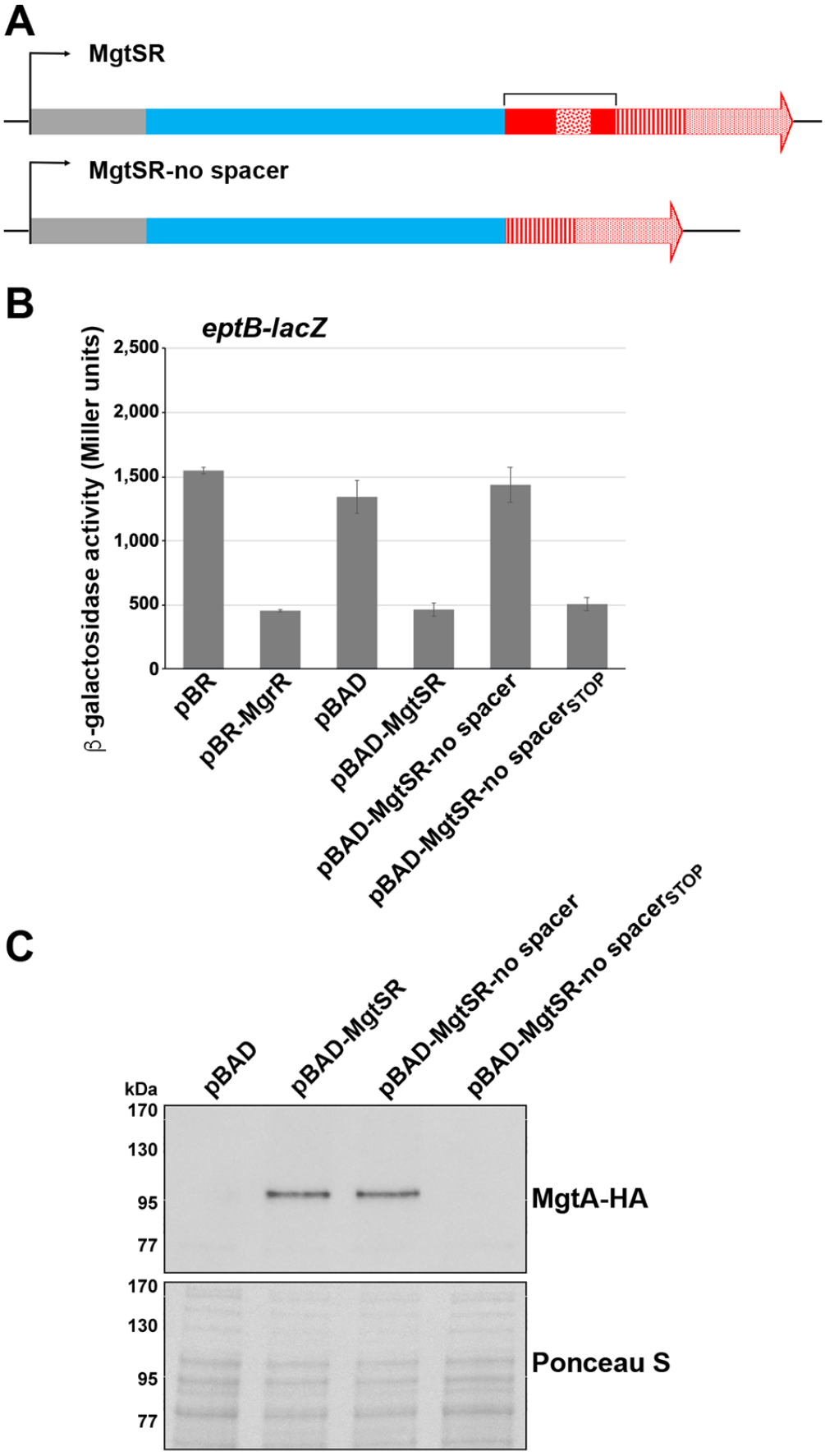
MgtS translation is the dominant activity absent the spacer sequence. (A) Schematic of the hybrid dual-function RNA with the spacer sequence (bracket) removed. Colors and patterns for plasmid, *mgtS*, *mgrR*, ARN, seed and terminator sequences are as in Fig. 1B. (B) Effect of pBR, pBR-MgrR, pBAD, pBAD-MgtSR, pBAD-MgtSR-no spacer, or pBAD-MgtSR_BP_ overexpression on an *eptB-lacZ* fusion in a Δ*mgrR* background. Graphs show average of three biological replicates with error bars representing standard deviation. (C) Immunoblot analysis of MgtA-HA cells in a Δ*mgtS* transformed with pBAD, pBAD-MgtSR, pBAD-MgtSR-no spacer, or pBAD-MgtSR-no spacer_STOP_ grown to OD_600_ 0.4 in N-media supplemented with 500 μM Mg^2+^, washed, and then resuspended and grown in N-media supplemented with arabinose but without Mg^2+^. Samples were collected after 20 min, and MgtA-HA was visualized with α-HA antibodies to detect MgtA-HA. The Ponceau S provides the loading control.

### Truncation of the spacer sequence differentially disrupts regulation by each component

To better understand how the composition of the 30 nt spacer sequence contributes to regulation by each component, we next examined the effects of truncations from either end of the sequence to determine if the 5′ or 3′ end played a more prominent role in maintaining the functions of the synthetic dual-function RNA. We thus constructed derivatives with five or ten nt truncations from either the 5′ or 3′ ends of the spacer sequence to give pBAD-MgtSR-5′Δ5nt, pBAD-MgtSR-5′Δ10nt, pBAD-MgtSR-3′Δ5nt, and pBAD-MgtSR-3′Δ10nt (Fig. 3A). The levels of these MgtSR deletion derivatives are similar to the full length MgtSR RNA, except for MgtSR-3′Δ10nt, which is somewhat lower (see Fig. S2A in supplemental material). We carried out β-galactosidase assays for *eptB-lacZ* strains carrying each of these plasmids or the control pBAD and pBAD-MgtSR plasmids. Interestingly, the derivatives with truncations near the 3′ end of the 30 nt sequence completely lost the ability to regulate *eptB-lacZ*, while the constructs with truncations near the 5′ end retained some ability to repress *eptB-lacZ* (Fig. 3B). When stop codons were introduced into each of the truncation constructs, the RNAs all repressed *eptB-lacZ* demonstrating that the truncated constructs retain the potential to base pair and translation of the sORF is responsible for disrupting regulatory activity.

**FIG. 3.**
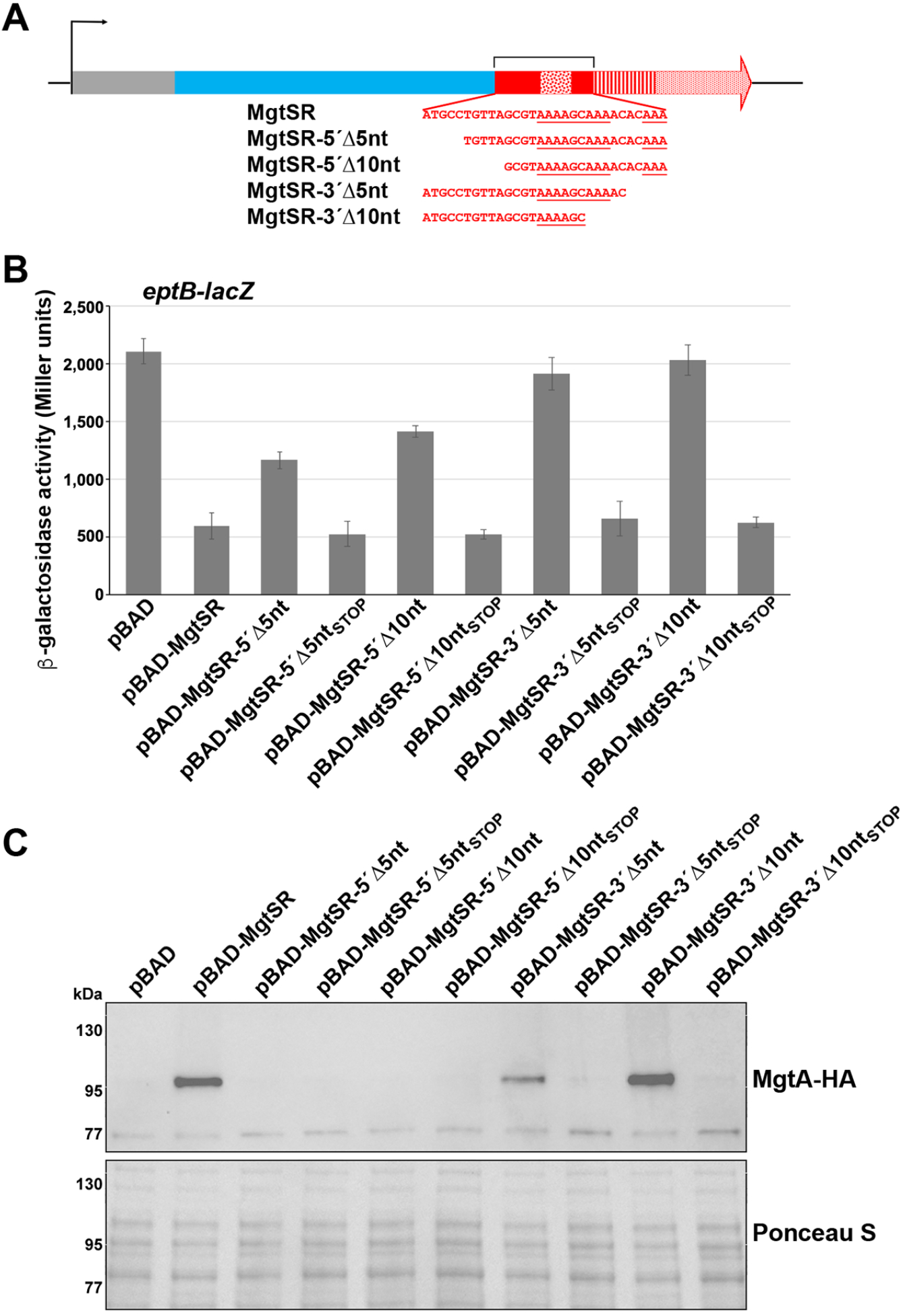
Analysis of the regulatory contributions of the spacer sequence. (A) Schematic of the 5-nt and 10-nt truncation mutants of MgtSR. Colors and patterns for plasmid, *mgtS*, *mgrR*, ARN, seed and terminator sequences are as in Fig. 1B. (B) Effects of overexpression from pBAD, pBAD-MgtSR, pBAD-MgtSR truncations, or pBAD-MgtSR truncation stop constructs on an *eptB-lacZ* fusion in a Δ*mgrR* background. Graphs show average of three biological replicates with error bars representing standard deviation. (C) Immunoblot analysis of MgtA-HA cells in a Δ*mgtS* transformed either pBAD, pBAD-MgtSR, pBAD-MgtSR truncations, or pBAD-MgtSR truncation stop constructs grown to OD_600_ 0.4 in N-media supplemented with 500 μM Mg^2+^, washed, and then resuspended and grown in N-media supplemented with arabinose but without Mg^2+^. Samples were collected after 20 min, and MgtA-HA was visualized with α-HA antibodies to detect MgtA-HA. The Ponceau S provides the loading control.

In contrast, the truncations towards the 5′ end of the spacer sequence interfered with the MgtSR effect on MgtA-HA levels, while the 3′ truncation derivatives still were capable of increasing MgtA-HA levels similar to MgtSR (Fig. 3C). Thus, while removal of portions of the spacer sequence from the 5′ end had less of an effect on base pairing activity, these deletions prevented MgtS-mediated stabilization of MgtA-HA. In contrast, the 3′ end truncation derivatives which lacked base pairing activity still stabilized MgtA-HA indicating MgtS function. It is possible that Hfq binding proximal to the ORF in the 5′Δ5nt and 5′Δ10nt derivatives has a negative effect on translation by competing with ribosome binding. Another possibility is that the truncations at either the 5′ or 3′ end produce changes in RNA structure that disrupt translation or base pairing, respectively. These results show that the specific sequence between the ORF and seed region has an impact on the functions of the chimeric RNA.

### The spacer sequence of the reverse MgtRS also impacts activity

To test whether the order of the encoded functions of the synthetic dual-function RNA impacted the activities, we placed the *mgtS* 5′ UTR including the RBS (denoted spacer sequence) and the *mgtS* coding sequence between the base pairing region and terminator sequence of *mgrR* (Fig. 4A). However, when cells with an *eptB-lacZ* fusion were transformed with pBAD and pBAD-MgtRS, we did not observe decreased β-galactosidase activity as for MgrSR (Fig. 4B). Furthermore, pBAD-MgtRS did not increase MgtA-HA levels as seen for pBAD-MgtSR (Fig. 4C). Northern analysis revealed that MgtRS was not detected indicating that the transcript might be unstable (see Fig. S2B in supplemental material). We hypothesized that the process of reversing the order of *mgtS* and *mgrR* coding elements might have disrupted binding to the Hfq protein if the ARN motif of MgrR is too distant from the terminator to allow for strong Hfq binding. Alternatively, the combinations of elements may have inadvertantly introduced an endonuclease cleavage site.

**FIG. 4.**
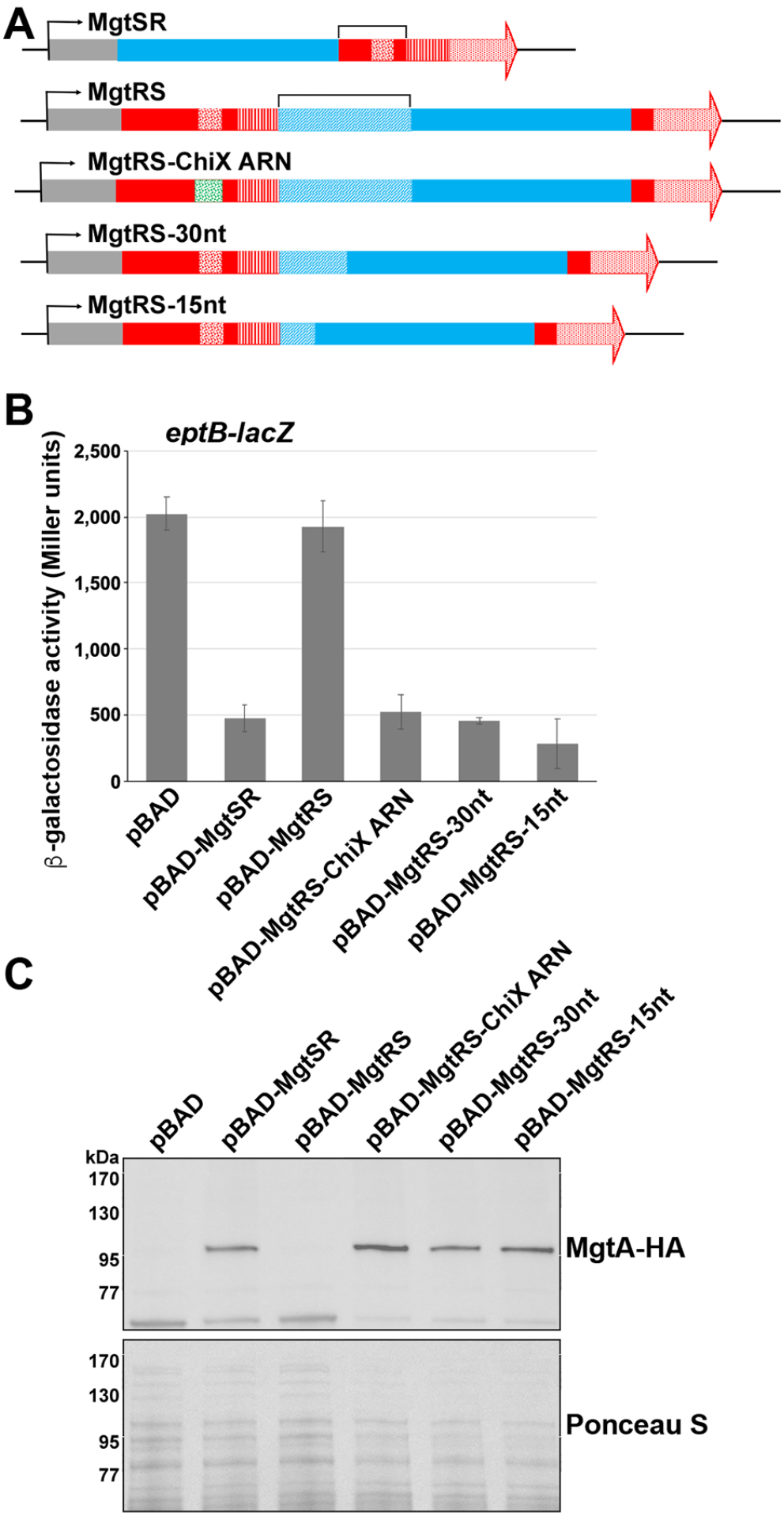
Reversal of components produces a nonviable construct in the absence of an appropriate spacer. (A) Schematic of the inverse hybrid constructs. Colors and patterns for plasmid, *mgtS*, *mgrR*, ARN, seed and terminator sequences are as in Fig. 1B. The stippled blue region corresponds to *mgtS* sequence outside of the coding region. (B) Effects of pBAD, pBAD-MgrSR, pBAD-MgtRS, pBAD-MgtRS-ChiX ARN, pBAD-MgtRS-30nt or pBAD-MgtRS-15nt overexpression on a *eptB-lacZ* fusion in a Δ*mgrR* background. Graphs show average of three biological replicates with error bars representing standard deviation. (C) Immunoblot analysis of MgtA-HA cells in a Δ*mgtS* transformed with pBAD, pBAD-MgrSR, pBAD-MgtRS, pBAD-MgtRS-ChiX ARN, pBAD-MgtRS-30nt, or pBAD-MgtRS-15nt grown to OD_600_ 0.4 in N-media supplemented with 500 μM Mg^2+^, washed, and then resuspended and grown in N-media supplemented with arabinose but without Mg^2+^. Samples were collected after 20 min, and MgtA-HA was visualized with α-HA antibodies to detect MgtA-HA. The Ponceau S provides the loading control.

**FIG 5.**
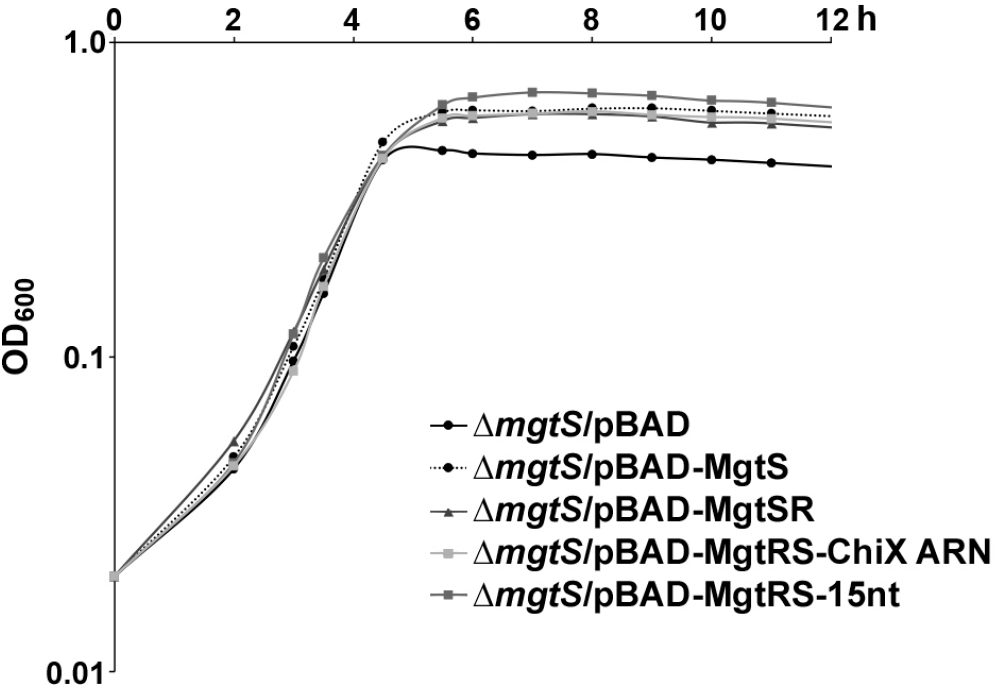
MgtSR and MgtRS constructs allow growth in low magnesium. Δ*mgtS::kan* strain (GSO229) transformed with pBAD, pBAD-MgtS, pBAD-MgtSR, pBAD-MgtRS-ChiX, or pBAD-MgtRS-15nt grown overnight in LB were diluted to OD_600_∼0.02 (time 0) in 25 ml of the N media with no Mg^2+^ and grown at 37°C. Growth was tracked by OD_600_ over 12h.

To attempt to stabilize the MgtRS derivative and gain activity, we made a number of modifications. First, we replaced the ARN sequence of *mgrR* in MgtRS with the strong ARN sequence from ChiX (MgtRS-ChiX ARN). Additionally, we made two other variants of MgtRS in which the 58 nt spacer sequence was truncated to 30 nt or 15 nt. We chose these lengths given the 30 nt spacer sequence in the functional MgtSR construct and the 15 nt spacer sequence in SgrS. When *eptB-lacZ* cells carrying pBAD-MgtRS-ChiX ARN, pBAD-MgtRS-30nt, and pBAD-MgtRS-15nt were assayed for β-galactosidase activity, we observed that the altered versions of MgtRS were all capable of *eptB-lacZ* repression (Fig. 4B). Additionally, pBAD-MgtRS-ChiX ARN, pBAD-MgtRS-30nt, pBAD-MgtRS-15nt were capable of increasing MgtA-HA levels to a similar degree as MgtSR (Fig. 4C). Northern analysis showed that all of the constructs could be detected, though we also observed some shorter products and MgtRS-15nt was present at elevated levels (see Fig. S2B in supplemental material). Together, these results show that functional synthetic dual-function RNAs can be generated and that the two functional elements, protein coding and base pairing, can be in either order as long as the appropriate spacer sequence is present.

### Synthetic dual-function RNAs complement the Δ*mgtS* growth defect in low Mg^2+^

Finally, we examined the ability of the synthetic dual-function RNAs to complement the low Mg^2+^ sensitivity of a Δ*mgtS* strain. While a growth defect was observed in limited Mg^2+^ media for the Δ*mgtS* strains carrying the pBAD vector control, the pBAD-MgtSR, pBAD-MgtRS-ChiX ARN, and pBAD-MgtRS-15nt constructs all protected the Δ*mgtS* cells at low intracellular Mg^2+^ levels similar to pBAD-MgtS. These data demonstrate that synthetic dual-function RNAs produce biologically relevant regulatory effects.

## DISCUSSION

Dual-function RNAs are an intriguing yet largely unexplored class of RNA regulatory molecules. Given the underidentification of small proteins, these RNA transcripts are likely more prevalent than has currently been reported. To better understand the organization and competition between the components of a dual-function RNA, we designed and constructed synthetic dual-function RNAs from the individual genes encoding the MgrR sRNA and MgtS small protein (Fig. 1A). These hybrid RNAs are capable of regulating by base pairing and are translated to give functional MgtS protein. We used derivatives of these synthetic constructs to examine how the organization of the components affects the regulatory activities of these RNA molecules.

### Elements necessary for dual function

The MgtSR and MgtRS chimeras and their derivatives underscore what elements are necessary to construct a dual-function RNA. First, translation of the sORF likely requires an unobstructed ribosome binding site. Proper sRNA function requires an unobstructed seed sequence for base pairing along with Hfq binding sites including a Rho-independent terminator. How these elements are spaced relative to one another has an impact on the two activities. We suspect that when the ARN sequence for Hfq binding was moved far from the Hfq binding site at the Rho-dependent terminator in the MgtRS construct, Hfq was no longer able to stabilize the RNA. This was overcome by replacing the *mgrR* ARN sequence with the longer *chiX* ARN sequence (Fig. 4). The length of the sequence that genetically separates the components of dual-function RNA also is important for maintaining the independent regulatory activity of each component. Most deletions that reduced the sequence separating the sORF and the seed sequence in MgtSR, eliminated base pairing activity (Fig. 2 and 3). The observation that all of these derivatives regained base pairing activity upon the introduction of a stop codon suggests the main obstacle to base pairing is occlusion by ribosome binding although some of the deletions might also affect Hfq binding. For most of these deletion constructs, the MgtS activity predominated, possibly because this activity is encoded first. Interestingly, however, with a limited distance between the two genetic elements, base pairing can also interfere with translation as illustrated for the 5′Δ5 nt and 5′Δ10 nt derivatives.

In other dual-function RNA such as AzuCR, the overlap of the ORF and base pairing regions leads to competition between the activities (14). Mutations that prevent translation of the ORF have been found to enhance regulation by the sRNA component. In contrast, for SgrS, where the components are separated by 15 nt, the introduction of mutations to disrupt translation of the encoded small protein, SgrT, do not change base pairing regulation of mRNA targets by SgrS (17). Furthermore, mutants with disrupted base pairing actually led to an increase in SgrT levels (17). This is consistent with the model that the stability of SgrS RNA dictates that translation will occur after base pairing regulation is completed. Overall, it is expected that separation of the parts of a dual-function RNA by a spacer sequence gives each component more autonomy despite existing within the same RNA molecule.

### Evolution of dual-function RNAs

In thinking about the components of dual-function sRNAs, it is interesting to consider how these regulatory molecules evolve (26) and whether there are more barriers to evolution of base pairing or expression of a small protein as the second feature. Phylogenetic analyses of SgrST and the RNAIII dual-function RNA from *Staphylococcus aureus* suggest that evolution in both directions is possible. For example, all SgrST homologs across enteric species carry the highly conserved 13 nt seed sequence with complementary to the *ptsG* RNA near the 3’ end of the RNA, while some species with a SgrS homolog, such as *Yersinia*, lack a functional SgrT ORF suggesting that the sRNA served as the precursor RNA which ultimately developed an mRNA function. In contrast, the 26 aa δ-haemolysin encoded by RNAIII appears to be more broadly conserved than the base pairing regions indicating an mRNA evolved base pairing capability (27).

### Possibility of exploiting dual-function RNAs

Dual-function RNAs present an interesting opportunity for synthetic biology applications. As we have demonstrated in this work, the creation of an artificial dual-function RNA from existing sRNA and small protein components is possible and can be tuned to favor one or the other activity. Given that these molecules can regulate both the translation and activity of a protein, dual function RNAs can provide an effective means to rapidly alter both mRNA and protein activity levels and thus allow efficient control of a specific process. Furthermore, if the two activities regulate different pathways, a single dual-function RNA can provide a regulatory link between the two pathways. By modulating when each component is active, these molecules could be used to regulate two pathways under mutually exclusive conditions. Dual-function RNAs represent an opportunity to design transcripts that can effectively control gene expression using two different regulatory mechanisms. Ultimately, there is much to be explored about the regulatory potential of natural and synthetic dual-function RNAs.

## MATERIALS AND METHODS

### Bacterial strains and plasmid construction

Bacterial strains, plasmids, and oligonucleotides used in this study are listed Supplemental Tables S1-S3. *E. coli* strains are derivatives of wild-type MG1655 (F- lambda- *ilvG*- *rfb*-50 *rph*-1. All mutations and constructs were confirmed by sequencing.

### Bacterial growth

Cells were grown to the indicated OD_600_ in Luria-Bertani broth (LB) overnight or N-minimal media (7.5 mM (NH_4_)_2_SO_4_, 0.5 mM K_2_SO_4_, 1 mM KH_2_PO_4_, 0.4% glycerol, 5 mM KCl, 100 mM Tris-HCl (pH 7.5), 10 μM Vitamin B1, and 0.1% casamino acids) with either 500 μM or 0 μM Mg^2+^ (28). pBAD24 and pBR322 plasmid constructs were induced with 0.4% arabinose or 1 mM IPTG. Where indicated, media contained antibiotics with the following concentrations: ampicillin (100 μg/ml), chloramphenicol (25 μg/ml) and kanamycin (30 μg/ml).

### β-galactosidase assays

Cultures were grown in LB to OD_600_∼1.0 with arabinose (0.2%). Cells (100 μl) were added to 700 μl of Z buffer (60 mM Na_2_HPO_4_•7H_2_O, 40 mM NaH_2_PO_4_•H_2_O, 10 mM KCl, 1 mM MgSO_4_ •7H_2_O, 50 mM β-mercaptoethanol). After adding 15 μl of freshly prepared 0.1% SDS and 30 μl of chloroform to each sample, the cells were vortexed for 30 s and then incubated at room temperature for 15 min to lyse the cells. The assay was initiated by the addition of 100 μl of ONPG (4 mg/ml) to each sample in 10 s intervals. The samples were incubated at room temperature before the reactions were terminated by the addition of 500 μl of 1M Na_2_CO_3_, after which A_420_ and A_550_ values were determined with a spectrophotometer and the absorbance data was used to calculate Miller units.

### Immunoblot analysis

After growth in the indicated medium cells corresponding to 0.5 OD_600_ were collected by centrifugation and rsuspended in 1X PBS (KD Medical), 7 μl of 2X Laemmli buffer (BioRad) and 2 μl of β-mercaptoethanol, and 10 μl were loaded on a Mini-PROTEAN TGX 5%–20% Tris-Glycine gel (Bio-Rad) and run in 1X Tris Glycine-SDS (KD Medical) buffer. The proteins were electro-transferred to nitrocellulose membranes (Invitrogen) for 1 h at 100 V. Membranes were blocked with 5% non-fat milk (BioRad) in 1X PBS with 0.1% of Tween 20 (PBS-T) for 1 h and probed with a 1:1,000 dilution of α-HA antiserum (Abcam) in the same PBS-T buffer with 5% milk for 1 h. After the incubation with the α-HA antiserum, membranes were incubated with a 1:20,000 dilution of HRP-labelled goat anti-rabbit antibody (Pierce). All blots were washed 4X with PBS-T and then developed with a Amersham ECL Western Blotting Detection Kit (GE Healthcare).

### Total RNA isolation

Cells corresponding to the equivalent of 10 OD_600_ were collected by centrifugation, and snap frozen in liquid nitrogen. RNA was extracted according to the standard TRIzol (Thermo Fisher Scientific) protocol. Briefly, 1 ml of room temperature TRIzol was add to cell pellets, resuspended thoroughly to homogenization, and incubated for 5 min at room temperature. After the addition of 200 μl of chloroform and thorough mixing by inversion, samples were incubated for 10 min at room temperature. After samples were centrifuged for 10 min at 4°C at 20000 x g, the upper phase (∼0.6 ml) was transferred into a new tube and 500 μl of isopropanol was added. Samples again were mixed thoroughly by inversion, incubated for 10 min at room temperature and centrifuged at 20000 x g for 15 min at 4°C. RNA pellets were washed twice with 75% ethanol and then dried at room temperature. RNA was resuspended in 20-50 μl of DEPC water and quantified using a NanoDrop (Thermo Fisher Scientific).

### Northern analysis

Total RNA (5-10 μg per lane) was separated on denaturing 8% polyacrylamide gels containing 6 M urea (1:4 mix of Ureagel Complete to Ureagel-8 (National Diagnostics) with 0.08% ammonium persulfate) in 1X TBE buffer at 300 V for 90 min. The RNA was transferred to a Zeta-Probe GT membrane (Bio-Rad) at 20 V for 16 h in 0.5X TBE, UV-crosslinked, and probed with ^32^P-labeled oligonucleotides (Listed in Table S1) in ULTRAhyb-Oligo buffer (Ambion Inc.) at 45°C. Membranes were rinsed twice with 2X SSC/0.1% SDS at room temperature, once with 0.2X SSC/0.1% SDS at room temperature, washed for 25 min with 0.2 × SSC/0.1% SDS at 45°C, followed by a final rinse with 0.2X SSC/0.1% SDS at room temperature before autoradiography was performed with HyBlot CL film (Denville Scientific Inc.).

### Growth curves

Colonies of Δ*mgtS::kan* (GSO229) transformed with pBAD, pBAD-MgtS, pBAD-MgtSR, pBAD-MgtRS-ChiX, and pBAD-MgtRS-15nt grown overnight in LB were diluted to OD_600_∼0.02 (time 0) in 25 ml of the N media with no Mg^2+^ and grown at 37°C. Growth was followed for 22 h.

## ACKNOWLEDGMENTS

We thank members of the Storz group for comments on the manuscript. This research was supported by the Intramural Research Program of the *Eunice Kennedy Shriver* National Institute of Child Health and Human Development.

## SUPPLEMENTAL FIGURE LEGENDS

**FIG. S1** Synthetic dual-function RNA constructs. List of MgtSR (sORF upstream of base pairing region) and MgtRS (base pairing region upstream of sORF) constructs. Bold black font indicates plasmid sequence, blue font indicates *mgtS* mRNA sequence, bold blue fonts indicates the *mgtS* ORF, black font indicates *mgrR* sequence, yellow indicates *mgrR* ARN motif, green indicates ChiX ARN motif, red indicates the terminator stem, *mgrR* base pairing region is underlined, and mutations are in bold red.

**FIG. S2** Levels of transcripts expressed from dual-function RNA constructs. (A) Northern analysis of Δ*mgtS* cells transformed with pBAD, pBAD-MgtSR, pBAD-MgtSR-no spacer, pBAD-MgtSR-no spacer_STOP_, pBAD-MgtSR-5′Δ5nt, pBAD-MgtSR-5′Δ10nt, pBAD-MgtSR-3′Δ5nt, or pBAD-MgtSR-3′Δ10nt. (B) Northern analysis of Δ*mgtS* cells transformed with pBAD, pBAD-MgtSR, pBAD-MgtRS, pBAD-MgtRS-ChiX ARN, pBAD-MgtRS-30nt, pBAD-MgtRS-15nt, or pBAD-MgtRS-stop. For both (A) and (B), Cells were grown in LB to OD∼1.0. Total RNA isolated was separated on an acrylamide gel, transferred to a membrane and sequentially probed for the 3′ region of *mgtS* and 5S (as a loading control).

